# The Neuroprotective Effects of *Ocimum gratissimum-*Supplemented Diet on Scopolamine-Induced Memory Impairment in Mice Model of Alzheimer’s Disease

**DOI:** 10.1101/2025.10.28.684872

**Authors:** Damilare Alabi Akanbi, Patrick Olarewaju Ogboriga, Ismaheel Akinwale Adeniyi, Akintunde Muthoir Lawal, Michael Ayomikun Olusola, Precious Ayomiposi Oluyemi, Samuel Adetunji Onasanwo

**Author notes:** Corresponding Author: Name: Samuel Adetunji Onasanwo, Address: University of Ibadan, Agbowo, Ibadan, Oyo state, Nigeria.

## Abstract

**Background:** Cognitive decline is a hallmark of Alzheimer’s disease, a progressive neurodegenerative illness primarily caused by the buildup of amyloid plaque, which is brought on by oxidative stress and neuroinflammation. Currently, therapeutic agents are focused on addressing clinical symptoms with associated side effects. This study aims to explore the neuroprotective potential of *Ocimum gratissimum,* a plant known for its richness in bioactive compounds. By investigating its effects on cognitive health, this study addresses a significant gap in the literature regarding dietary interventions for Alzheimer’s disease.

**Methods:** Thirty-six animals were divided into six groups of six mice each: the control group received distilled water intraperitoneally; the scopolamine group received only scopolamine (1 mg/kg i.p.); the three test groups were fed 5%, 10%, and 20% *Ocimum gratissimum*-supplemented diets while also receiving scopolamine (1 mg/kg i.p.); and the positive control group received Donepezil (5 mg/kg) followed by the injection of scopolamine (1 mg/kg i.p.). Donepezil was administered orally 30 minutes before the scopolamine injection. All treatments were administered daily for 14 consecutive days.

**Results:** The supplemented diet groups showed significantly improved spatial memory and navigation compared to the scopolamine-only group. Biochemical analyses revealed that *O. gratissimum* mitigated scopolamine-induced oxidative stress and neuroinflammation, with marked improvements in antioxidant enzyme levels, reduced lipid peroxidation, and modulation of pro-inflammatory cytokines.

**Conclusion:** Dietary intervention using *Ocimum gratissimum* leaf was able to improve spatial memory and protect against memory impairment, suggesting its potential as a neuroprotective agent against Alzheimer’s disease-like pathology.

## INTRODUCTION

Alzheimer’s disease (AD) continues to be the most common type of dementia, accounting for approximately 60-70% of dementia cases worldwide (Alzheimer’s Association, 2023). Alzheimer’s usually starts slowly and gets worse over time, eventually causing significant impairments in language, reasoning, memory, and judgment (Veljkovic *et al*., 2018). It is a progressive neurodegenerative disorder that impairs everyday functioning, causes memory loss, and cognitive decline. Neuroinflammation, synaptic dysfunction, and the accumulation of tau protein tangles and amyloid beta plaques in the brain are hallmarks of AD pathology (Tenchov *et al*., 2024). Alzheimer’s disease (AD) is becoming more and more common worldwide, which emphasizes how urgently effective treatments are needed. There is no solid proof that cholinesterase inhibitors like donepezil or NMDA receptor antagonists (memantine) alter the underlying disease process in Alzheimer’s or any other form of dementia, even though they offer some short-term symptom relief (NICE, 2018). It is evident that current treatments do not change the course of the disease, and the lack of progress in therapeutic advancements highlights the need for better alternatives.

The two main contributors to Alzheimer’s disease (AD) progression are oxidative stress and neuroinflammation. An imbalance between the generation of reactive oxygen species (ROS) and the antioxidant defense system of the cell causes oxidative stress, which results in major biochemical changes that cause neurodegeneration (Chen *et al*., 2012). Chronic neuroinflammation has become a key component of AD pathogenesis, serving as a primary driver of disease progression rather than just a byproduct (Toader *et al*., 2024). The primary immune cells in the central nervous system, microglia and astrocytes, are essential to this process. They are unquestionably important in the development of AD because of their continuous activation, which causes the release of pro-inflammatory mediators that exacerbate oxidative stress and cause synaptic dysfunction (Colonna *et al*., 2021; Li *et al*., 2024; Yang *et al*., 2025).

Scopolamine is a muscarinic receptor antagonist that blocks cholinergic neurotransmission, leading to memory impairment in Rodents. Recent studies have reported that Scopolamine increases the accumulation of reactive oxygen species, which induces oxidative stress, leading to memory impairment (Bunadri *et al*., 2013a). Administering scopolamine has been demonstrated to cause deficits in spatial navigation and impaired cognitive abilities like learning and memory, which are similar to the symptoms seen in Alzheimer’s patients (Lohninger *et al*., 2015). Because of this, scopolamine is a useful psychopharmacological model for Alzheimer’s (Bunadri *et al*., 2013b). Scopolamine has also been connected to increased neuroinflammation, which is a contributing factor to cognitive impairments (Cheon *et al*., 2021). The mechanisms of neuroinflammation and oxidative stress work together to highlight the value of the scopolamine model in researching the course of memory impairment as seen in Alzheimer’s disease.

Natural products have become attractive therapeutic substitutes, especially because of their low toxicity profiles and multitarget mechanisms. Due to its varied phytochemical makeup, *Ocimum gratissimum*, a plant that belongs to the Lamiaceae family and has been used for ethnomedical purposes throughout Africa and Asia for a long time, stands out among these. It is abundant in phenolic compounds, flavonoids, eugenol, and thymol, all of which have been demonstrated to have potent neuroprotective, anti-inflammatory, and antioxidant qualities (Akara *et al*., 2021). According to studies, extracts from *O. gratissimum* can suppress lipid peroxidation, scavenge free radicals, and alter immune responses (Njan *et al*., 2023). Although traditional medicine uses it extensively to treat ailments like fever, diabetes, and respiratory infections, its potential in neurodegenerative research, particularly in AD, is mainly unknown.

Limited research has indicated that it may have CNS-modulating properties, including sedative and anxiolytic effects (Okonkwo *et al*., 2017). However, research on the impact of *O. gratissimum* dietary supplementation on memory function, and neurochemical indices in a model similar to AD is still lacking. Moreover, no research has examined its impact on nitrosative stress, cholinergic enzyme activity, oxidative stress markers, and cytokine levels all at once within the same experimental setup.

A scopolamine-induced mouse model of Alzheimer’s disease was used to assess the neuroprotective effectiveness of a diet supplemented with *Ocimum gratissimum*. To evaluate memory and anxiety-related behaviors, a variety of behavioral paradigms were employed, such as the Elevated Plus Maze (EPM) and Novel Object Recognition (NOR). Important markers of neuroprotection, inflammation, and oxidative stress were measured by thorough biochemical analyses. By combining behavioral and biochemical endpoints, this study offers new proof that *Ocimum gratissimum* may reduce AD-related cognitive decline and neuropathological alterations, which could help direct the creation of dietary-based treatment approaches for neurodegenerative illnesses.

## MATERIALS AND METHODS

### Drugs and Reagents

Scopolamine, *Ocimum gratissimum*, ELISA kits for Tumor Necrotic Factor-alpha (TNF-alpha) and interleukin-6 (IL-6), Donepezil, Phosphate Buffered Solution, Normal Saline, Formaldehyde, Methylated Spirit. The other chemicals and reagents used were of analytical grade.

### Plant Material

Fresh leaves of *Ocimum gratissimum* were collected from a local farm in Ibadan, Oyo State, Nigeria. The plant was identified and authenticated at the Department of Botany, University of Ibadan, and a voucher specimen was deposited at the university herbarium under the number UIH/OG/024. The leaves were thoroughly rinsed with clean water, air-dried under shade at room temperature for seven days, and then ground into a fine powder using an electric blender. The powdered Ocimum gratissimum was incorporated into the mice diet at the appropriate concentrations to prepare the supplemented diet used in the study.

### Drugs and Chemicals

Scopolamine hydrobromide (≥98% purity) was purchased from Sigma-Aldrich (St. Louis, MO, USA). All other chemicals and reagents used in the biochemical assays, including those for the estimation of lipid peroxidation, reduced glutathione, nitrite, catalase, superoxide dismutase, glutathione S-transferase, acetylcholinesterase activity, and pro-inflammatory cytokines (TNF-α and IL-6), were of analytical grade and obtained from Sigma-Aldrich and Randox Laboratories Ltd (UK), unless otherwise stated.

### Feed Supplementation Process

Diets were made from commercially formulated feed (KesMac Feeds, Ibadan, Nigeria) containing crude protein 23.00%, fat/oil 6.00%, crude fiber 5.00%, calcium 1.00%, available phosphorus 0.40%, lysine 1.20%, methionine 0.50%, salt (min) 0.30%, and metabolizable energy (min) 2900 kcal/kg. The *Ocimum gratissimum*-supplemented diet was formulated to contain 5%, 10%, and 20% *Ocimum gratissimum*. 5% *Ocimum gratissimum* supplemented diet has 0.25kg of the *Ocimum gratissimum* sample mixed with 4.75kg of standard diet, while 0.5kg and 1kg of the sample were mixed with 4.5kg and 4kg of the standard diet for 10% and 20% supplementation, respectively. Supplemented and non-supplemented diets were homogenized using a mixer and made into pellet (Anim, 2023).

### Preparation of Scopolamine

Scopolamine stock solution was prepared, with a serial dose of 1mg/kg.

### Preparation of Donepezil

Donepezil was selected for this study as a non-steroidal anti-inflammatory drug comparator of the effectiveness of *Ocimum gratissimum* treatment against neuroinflammation. The stock solution was prepared, a serial dose of 5mg/kg (Xiao, 2018).

### Route of Administration

*Ocimum gratissimum* was supplemented with the diet, while Scopolamine and Donepezil were administered intraperitoneally. All treatment procedures were conducted in the morning.

### Animals

Thirty-six male Swiss mice with 25-35g body weight were randomly selected for this study. After separating them according to their weights, cages for the study were labeled according to the experimental design. A technician, blinded to the housing labels, was instructed to place animals at random into the labeled cages, and treatments were administered according to these labels.

The animals were obtained from the University of Ibadan Department of Physiology’s postgraduate animal house and housed in plastic cages with wire meshed covers for proper ventilation in the same animal house. The animals were given access to water and rodent chow *ad libitum* and maintained under normal light and dark cycles (12h day/12h night), acclimatized. Experimental protocol and sample size followed the standard procedure following the Principle National Institution of Health, with ethical approval number NHREC/UI-ACUREC/05/12/2022A.

### Experimental Design

The animals were divided into six groups of six mice each: the control group received distilled water intraperitoneally; the scopolamine group received only scopolamine (1 mg/kg i.p.); the three test groups were fed 5%, 10%, and 20% *Ocimum gratissimum*-supplemented diets while also receiving scopolamine (1 mg/kg i.p.); and the positive control group received Donepezil (5 mg/kg) followed by the injection of scopolamine (1 mg/kg i.p.). Donepezil was administered orally 30 minutes before the scopolamine injection. All treatments were administered daily for 14 consecutive days.

### Behavioral Assessments

All animals used in this study were subjected to behavioral experiments with no exclusion, and behavioral protocols were done with the experimenter blinded to the testing groups to prevent bias.

### Novel Object Recognition

The experiment was carried out to assess the animal’s ability to discriminate between a familiar and a novel object. Briefly, the test consisted of a habituation phase, where the animals were allowed to freely explore an empty arena for 10 minutes. The test phase was conducted the following day during which the animals were placed back in the arena with one familiar and one novel object. The time spent exploring each object was recorded for 5 minutes using a stopwatch and a discrimination index was calculated to determine the animal’s preference for the novel object.

### Elevated Plus Maze

Anxiety-like behavior was assessed using the elevated plus maze (EPM) test. The EPM apparatus consisted of four arms (two open and two closed) elevated 50 cm from the ground. Each animal was placed in the center of the maze, facing one of the open arms, and allowed to freely explore the maze for 5 minutes. The time taken for the animal to move from the center of the maze to one of the closed arms (transfer latency) was recorded as a measure of anxiety-like behavior, with longer transfer latencies indicating greater anxiety (Walf & Frye, 2007).

### Tissue Preparation

The mice were sacrificed by cervical decapitation right after the behavioral tests, and their brains were promptly taken out and put in boxes with frozen saline to solidify for ten minutes. To create the 10% homogenate, the brain was placed inside a graduated cylinder and PBS (pH 7.4) was added. After centrifuging each tube for 15 minutes at 10,000 rpm/min at 4°C, the supernatant was taken out for biochemical analyses.

## BIOCHEMICAL ASSAYS

### Determination of Lipid peroxidation

Lipid peroxidation was evaluated by measuring the levels of malondialdehyde (MDA), a marker of oxidative stress, using the thiobarbituric acid reactive substances (TBARS) assay. Briefly, brain samples were homogenized in ice-cold phosphate buffer (pH 7.4) and centrifuged at 10,000 × g for 20 minutes at 4°C. The supernatant was collected, and protein concentration was determined using the bicinchoninic acid (BCA) assay (Smith *et al*., 1985). MDA levels were then measured by reacting the tissue supernatant with thiobarbituric acid and quantifying the pink chromogen spectrophotometrically at 532 nm. Malondialdehyde concentrations were expressed as nmol per mg of protein.

### Determination of Acetylcholinesterase Activity

Acetylcholinesterase (AChE) activity was measured in brain tissue homogenates as an indicator of cholinergic function. Briefly, brain samples were homogenized in ice-cold phosphate buffer (pH 8.0) and centrifuged at 10,000 × g for 20 minutes at 4°C. The supernatant was collected, and protein concentration was determined using the bicinchoninic acid (BCA) assay. Acetylcholinesterase activity was then measured by mixing the tissue supernatant with acetylthiocholine iodide as the substrate and 5,5’-dithiobis-(2-nitrobenzoic acid) (DTNB) as the chromogen, as described by (Ellman *et al*., 1961) with minor modifications (Nordberg *et al*., 1989). The rate of change in absorbance at 412 nm was monitored spectrophotometrically for 5 minutes, and AChE activity was calculated and expressed as μmol of acetylthiocholine hydrolyzed per minute per mg of protein.

### Determination of Glutathione Activity

Glutathione (GSH) levels were measured in brain homogenates as an indicator of antioxidant status (Forman *et al*., 2009). Briefly, brain samples were homogenized in ice-cold phosphate buffer (pH 7.4) containing EDTA, and the supernatant was collected after centrifugation at 10,000 rpm for 20 minutes at 4°C. GSH levels were determined by reacting the tissue supernatant with 5,5’-dithiobis (2-nitrobenzoic acid) (DTNB) and measuring the absorbance at 412 nm. GSH concentrations were expressed as μmol per mg of protein.

### Determination of Nitrite levels

Nitric oxide (NO) levels were indirectly assessed by measuring nitrite concentrations, a stable metabolite of NO, using the Griess reaction (Guevara *et al*., 1998; Tsikas, 2007). Brain samples were homogenized in ice-cold phosphate buffer (pH 7.4), and the supernatant was collected after centrifugation at 10,000 × g for 20 minutes at 4°C. The tissue supernatant was then incubated with the Griess reagent, and the absorbance was measured at 540 nm. Nitrite levels were expressed as μmol per mg of protein.

### Determination of Catalase Activity

Catalase activity was measured in brain homogenates as an antioxidant enzyme (Aebi, 1984; Goth, 1991). Briefly, brain samples were homogenized in ice-cold phosphate buffer (pH 7.0) and centrifuged at 10,000 × g for 20 minutes at 4°C. The supernatant was collected, and protein concentration was determined using the BCA assay. Catalase activity was then measured by monitoring the decomposition of hydrogen peroxide (H2O2) spectrophotometrically at 240 nm. Catalase activity was expressed as μmol of H2O2 consumed per minute per mg of protein.

### Determination of Superoxide Dismutase Activity

Superoxide dismutase (SOD) activity was measured in brain homogenates as an antioxidant enzyme (Misra *et al*., 1972; Nishikimi *et al*., 1972). Brain samples were homogenized in ice-cold phosphate buffer (pH 7.4) and centrifuged at 10,000 × g for 20 minutes at 4°C. The supernatant was collected, and protein concentration was determined using the BCA assay. SOD activity was then assessed by measuring the inhibition of the auto-oxidation of epinephrine, and the absorbance was read at 480 nm. SOD activity was expressed as units per mg of protein.

### Determination of Glutathione S-Transferase Activity

Glutathione S-transferase (GST) activity was measured in brain homogenates as a phase II detoxification enzyme (Habig *et al*., 1974; Sheehan *et al*., 2001). Brain samples were homogenized in ice-cold phosphate buffer (pH 6.5) and centrifuged at 10,000 × g for 20 minutes at 4°C. The supernatant was collected, and protein concentration was determined using the BCA assay. GST activity was then measured by monitoring the conjugation of glutathione with 1-chloro-2,4-dinitrobenzene (CDNB) spectrophotometrically at 340 nm. GST activity was expressed as μmol of CDNB-GSH conjugate formed per minute per mg of protein.

### Estimation of Proinflammatory Cytokines (TNF-α, IL-6)

Levels of the proinflammatory cytokines tumor necrosis factor-alpha (TNF-α) and interleukin-6 (IL-6) were measured in brain homogenates using commercially available enzyme-linked immunosorbent assay (ELISA) kits (Debnath *et al*., 2013). Brain samples were homogenized in ice-cold lysis buffer containing protease inhibitors, and the supernatant was collected after centrifugation at 10,000 × g for 20 minutes at 4°C. TNF-α and IL-6 levels were then quantified according to the manufacturers’ instructions, and the results were expressed as pg per mg of protein.

## STATISTICAL ANALYSIS

Data were analysed using GraphPad Prism software version 8.3.4. The data were presented as the mean ± SEM (n = 5). For normally distributed data, comparison among the different studied groups was done using one-way ANOVA, followed by the Tukey multiple comparison test. The Scopolamine group was analysed compared to the control group, and the groups receiving the extract were compared to the Scopolamine group. The difference was taken to be statistically significant at p < 0.05.

## RESULTS

### *Ocimum gratissimum*-supplemented diet ameliorated non-spatial memory impairment as induced by Scopolamine

The effect of the *Ocimum gratissimum*-supplemented diet on memory impairment as assessed by the Novel Object Recognition Test (NORT) is shown in Figure 1. The results indicate a significant decrease (p < 0.001) in the discrimination index of the Scopolamine group compared to the control group, with a value of -0.10 ± 0.04. In contrast, the groups receiving 5% and 10% *Ocimum gratissimum* showed discrimination indices of 0.45 ± 0.05 and 0.50 ± 0.06, respectively. The positive control group treated with Donepezil exhibited a discrimination index of 0.55 ± 0.07. These indices show the performance of the control group that received no treatment, which had a discrimination index of 0.60 ± 0.06.

**Figure 1.**
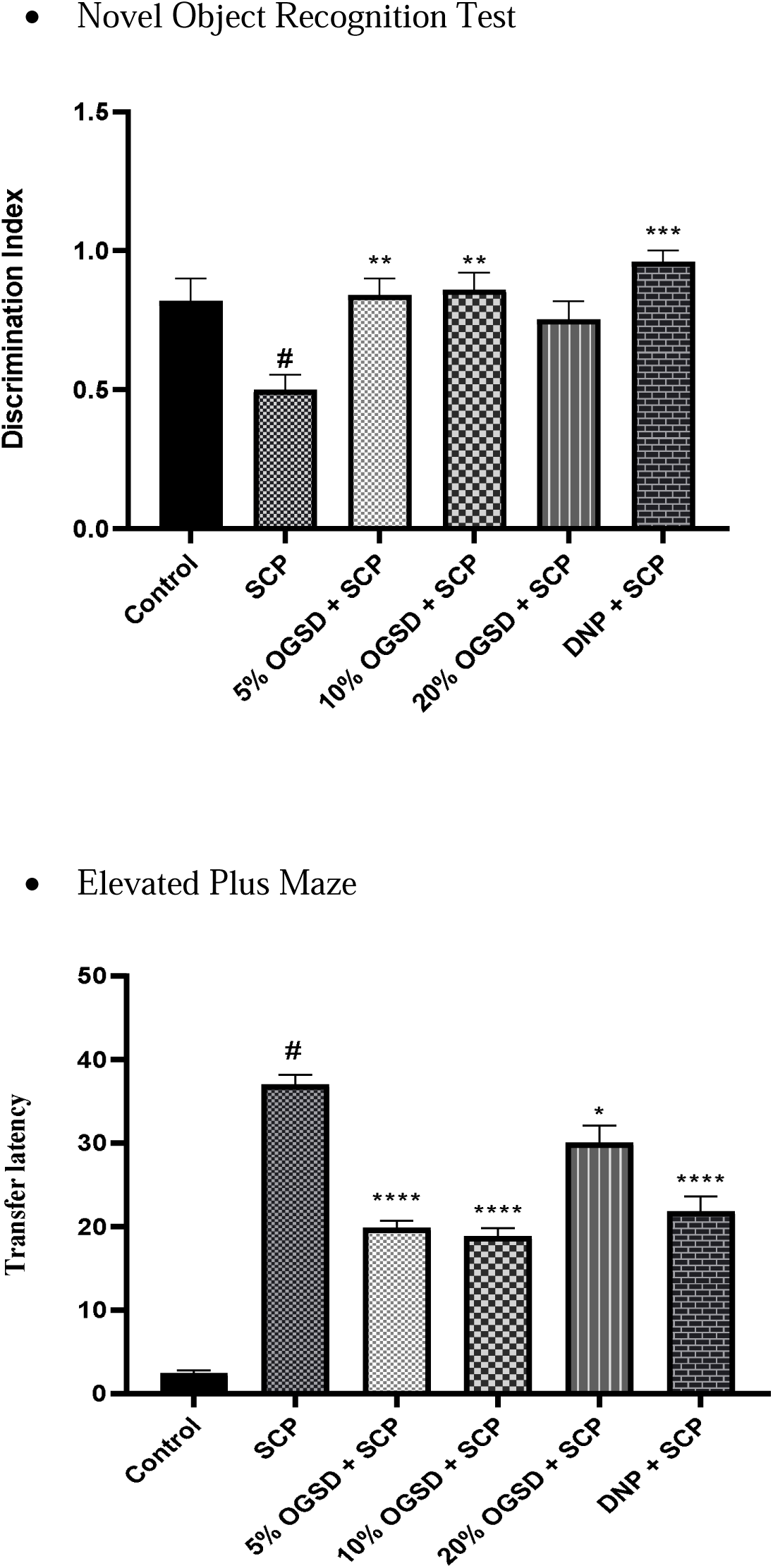
**A-B:** Effect of *Ocimum gratissimum*-supplemented diet on discrimination index and transfer latency in Scopolamine-induced memory impairment. Values are represented as mean ± SEM (n=5). The data were analyzed using a One-Way Analysis of Variance (ANOVA) followed by a Sidak’s posthoc test. The significance level is expressed as # p < 0.05 for Scopolamine vs Control, * p < 0.05 for *Ocimum gratissimum* vs Scopolamine. SCP is Scopolamine, OGSD is *Ocimum gratissimum*-supplemented diet and DNP is Donepezil.

### Effect of *Ocimum gratissimum*-supplemented diet on transfer latency in Scopolamine-induced memory impairment

The effect of *Ocimum gratissimum* on anxiety was assessed using the Elevated plus maze test and the data obtained were analyzed using a one-way ANOVA for the time spent in the open and closed arms of the apparatus.

The effect of *Ocimum gratissimum* on anxiety was assessed using the Elevated Plus Maze test, as shown in Figure 1B. The results indicate a significant increase (p < 0.001) in transfer latency in the Scopolamine group compared to the control group, with a value of 45.3 ± 2.1 seconds. The groups receiving 5% and 10% *Ocimum gratissimum* showed transfer latencies of 30.2 ± 1.5 seconds and 28.4 ± 1.3 seconds, respectively. The positive control group treated with Donepezil had a transfer latency of 26.5 ± 1.2 seconds. These latencies reflect the performance of the control group that received no treatment, which had a transfer latency of 24.1 ± 1.0 seconds.

### Effects of *Ocimum gratissimum*-supplemented diet on levels of MDA in the hippocampus and prefrontal cortex of scopolamine-induced memory impairment in Swiss mice

Malondialdehyde (MDA) levels were estimated in the hippocampus to assess lipid peroxidation, as shown in Figure 2A. The results indicate a significant increase in MDA levels in the Scopolamine group (3.71 ± 0.24) compared to the control group (F(5, 12) = 16.73; P < 0.0001). MDA was significantly reduced in the *Ocimum gratissimum*-supplemented diet (OGSD) groups, with levels of 2.24 ± 0.31, 2.01 ± 0.45, and 2.26 ± 0.48 for the 5%, 10%, and 20% diets, respectively. The positive control group treated with Donepezil showed MDA levels of 3.26 ± 0.09.

**Figure 2.**
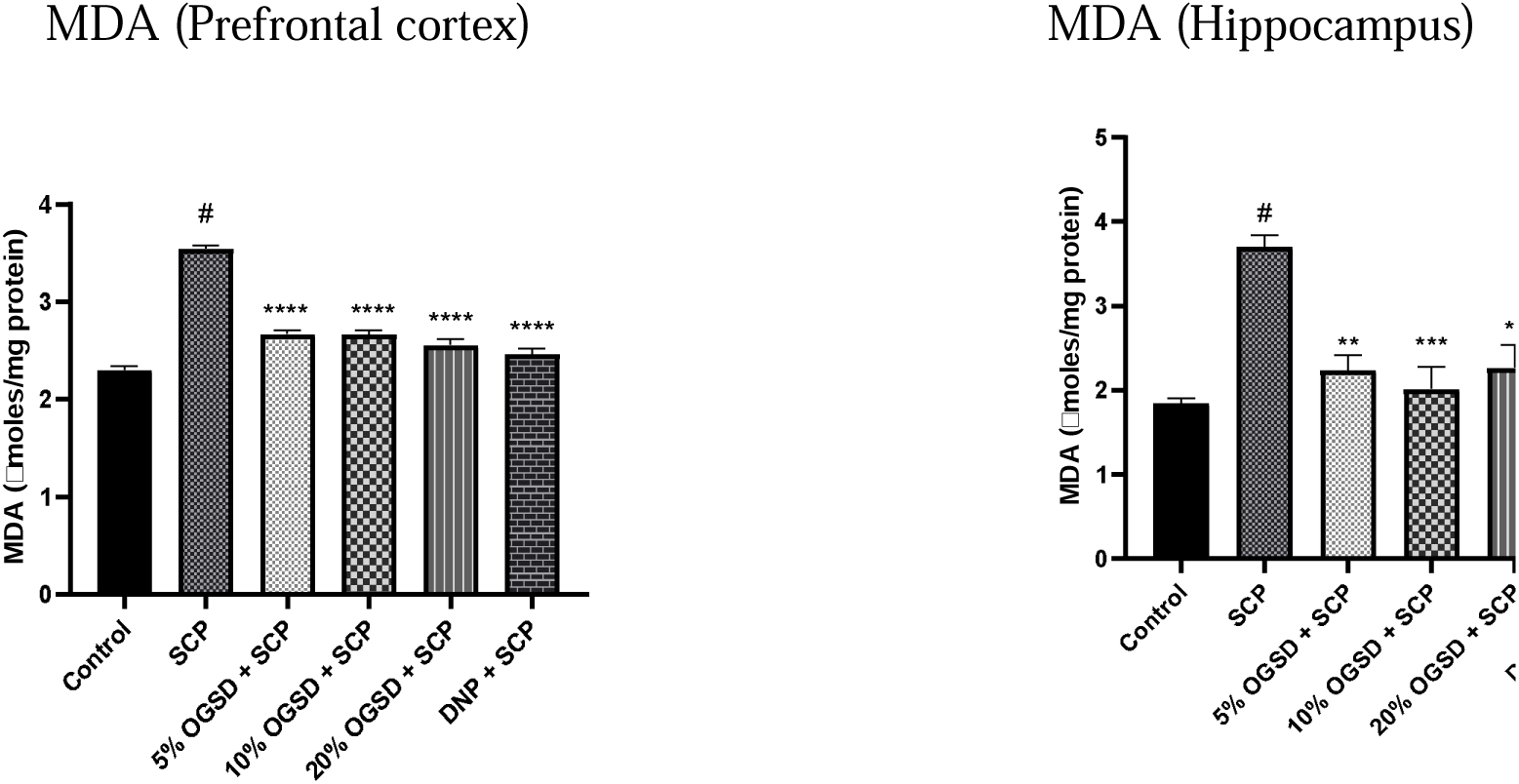
**A-B:** Effect of *Ocimum gratissimum*-supplemented diet on Malondialdehyde levels in Scopolamine-induced memory impairment. Values are represented as mean ± SEM (n=5). The data were analyzed using a One-Way Analysis of Variance (ANOVA) followed by a Sidak’s posthoc test. The significance level is expressed as # p < 0.05 for Scopolamine vs Control, * p < 0.05 for *Ocimum gratissimum* vs Scopolamine. SCP is Scopolamine, OGSD is *Ocimum gratissimum*-supplemented diet and DNP is Donepezil.

MDA levels were also estimated in the prefrontal cortex for further assessment of lipid peroxidation, as shown in Figure 2B. The Scopolamine group had elevated MDA levels of 3.55 ± 0.06 (F(5, 12) = 84.63; P < 0.0001). Significant reductions in MDA were observed in the OGSD groups, with values of 2.67 ± 0.07, 2.66 ± 0.08, and 2.55 ± 0.10, respectively. Donepezil treatment resulted in MDA levels of 2.46 ± 0.10.

### Effects of *Ocimum gratissimum* on levels of Glutathione on the hippocampus and prefrontal cortex of scopolamine-induced memory impairment in Swiss mice

As shown in Figure 3A, *Ocimum gratissimum* significantly increased the glutathione level in the hippocampus (F (5, 12) = 22.52; P<0.0001). Scopolamine administration significantly reduced glutathione level when compared to the control. Treatment with OGSD (5% and 20%) and Donepezil significantly increased glutathione level (3.50 ± 0.36, 3.46 ± 0.06 and 4.23 ± 0.21) when compared with the Scopolamine-only group (2.30 ± 0.51).

**Figure 3.**
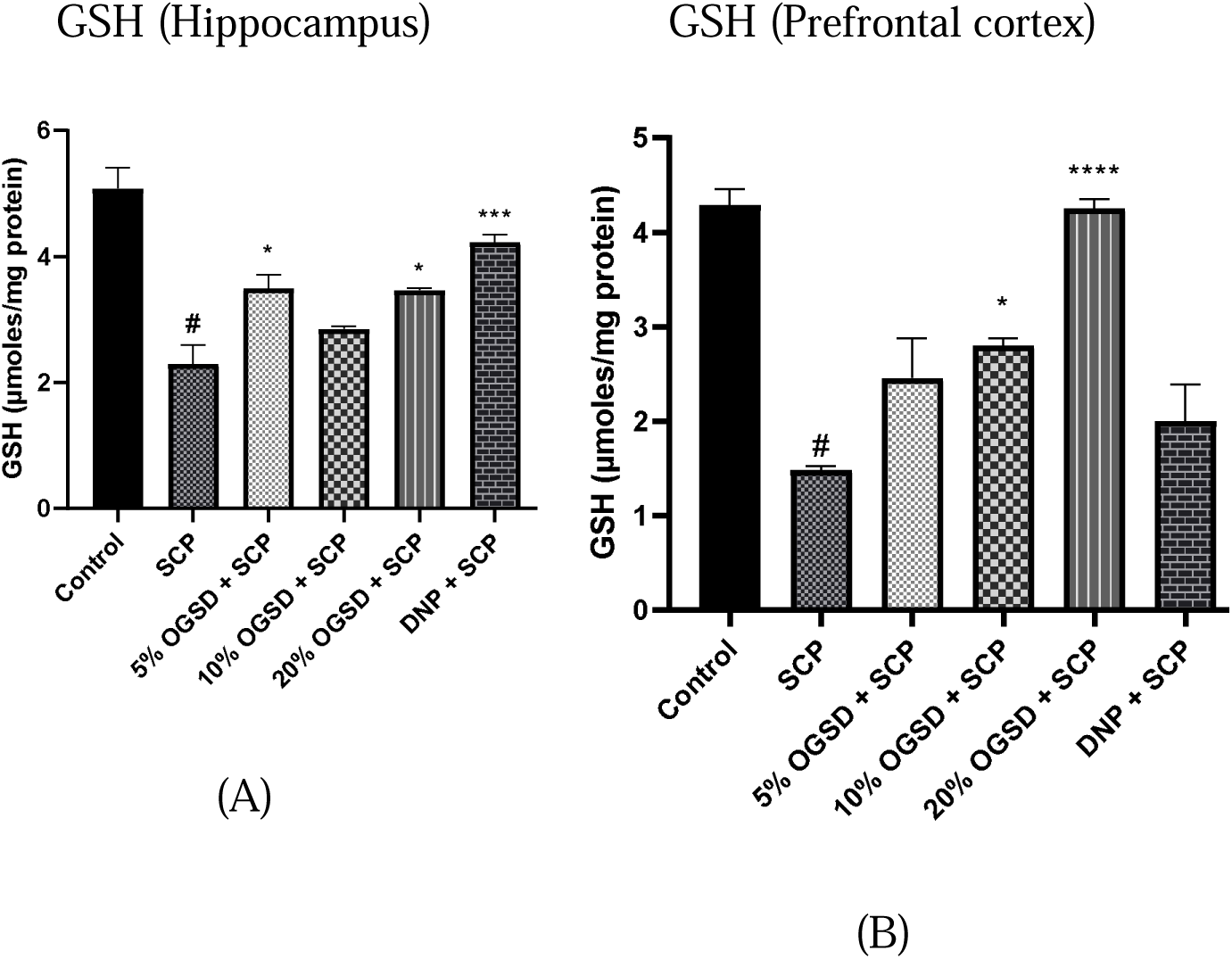
**A-B:** Effect of *Ocimum gratissimum*-supplemented diet on Glutathione levels in Scopolamine-induced memory impairment. Values are represented as mean ± SEM (n=5). The data were analyzed using a One-Way Analysis of Variance (ANOVA) followed by a Sidak’s posthoc test. The significance level is expressed as # p < 0.05 for Scopolamine vs Control, * p < 0.05 for *Ocimum gratissimum* vs Scopolamine. SCP is Scopolamine, OGSD is *Ocimum gratissimum*-supplemented diet and DNP is Donepezil.

As shown in Figure 3B, Scopolamine administration significantly reduced (1.49 ± 0.07) glutathione level in the prefrontal cortex (F (5, 12) = 21.96; P<0.0001) when compared to the control (4.29 ± 0.30). Treatment with OGSD (20%) significantly increased glutathione level (4.25 ± 0.17) when compared with the Scopolamine-only group (1.49 ± 0.07).

### Effects of *Ocimum gratissimum*-supplemented diet on Nitrite levels on the hippocampus and prefrontal cortex of scopolamine-induced memory impairment in Swiss mice

Nitrite levels in the hippocampus were significantly increased in the Scopolamine-only group (4.97 ± 0.16) compared to the control group (2.55 ± 0.28), as shown in Figure 4A (F(5, 12) = 30.90; P < 0.0001). Treatment with the *Ocimum gratissimum*-supplemented diet (OGSD) at 5%, 10%, and 20% and Donepezil resulted in reduced nitrite levels of 2.86 ± 0.24, 2.92 ± 0.02, 3.50 ± 0.27, and 3.69 ± 0.45, respectively, compared to the Scopolamine-only group.

**Figure 4.**
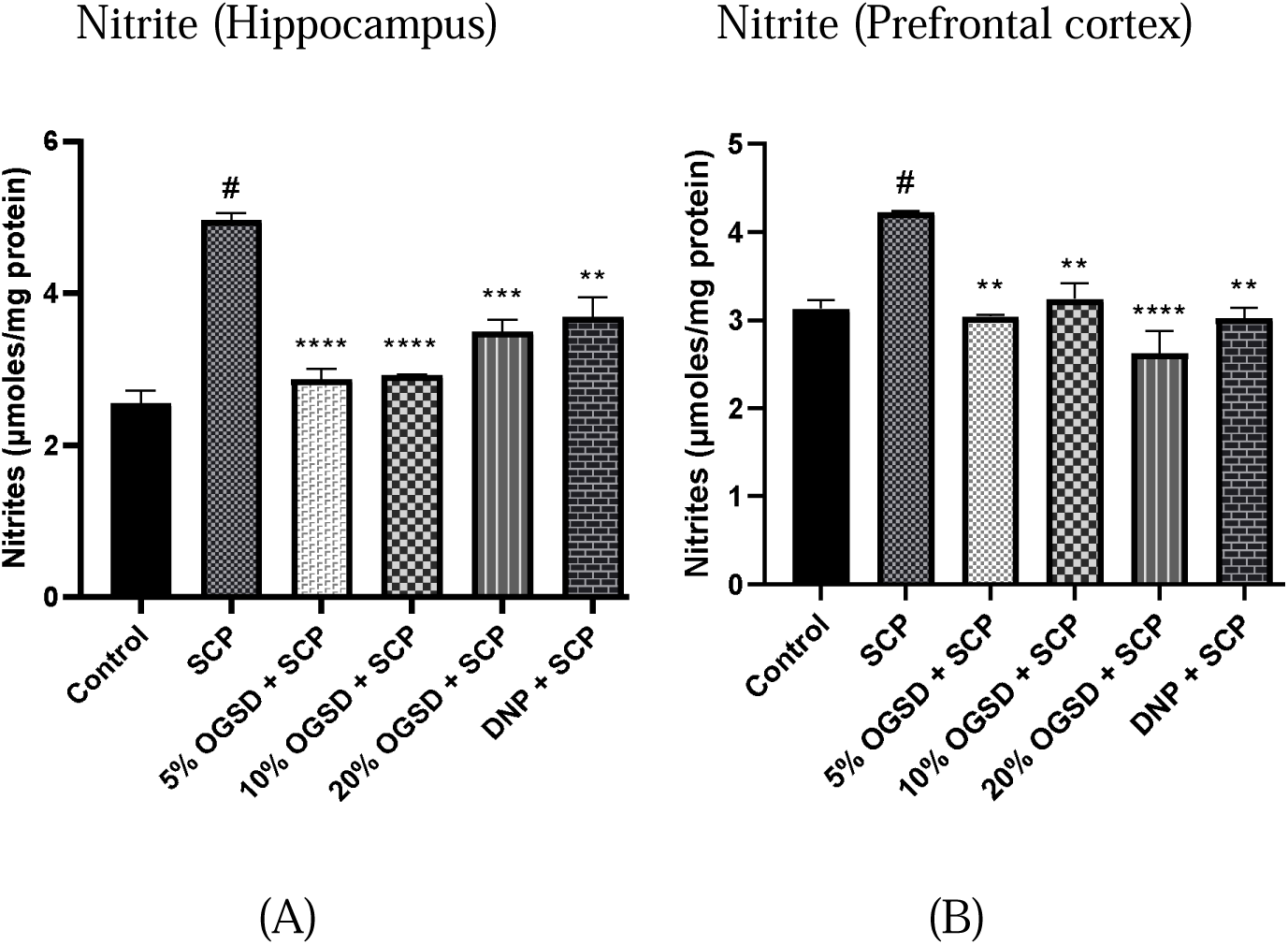
**A-B:** Effect of *Ocimum gratissimum*-supplemented diet on Nitrite levels in Scopolamine-induced memory impairment. Values are represented as mean ± SEM (n=5). The data were analyzed using a One-Way Analysis of Variance (ANOVA) followed by a Sidak’s posthoc test. The significance level is expressed as # p < 0.05 for Scopolamine vs Control, * p < 0.05 for *Ocimum gratissimum* vs Scopolamine. SCP is Scopolamine, OGSD is *Ocimum gratissimum*-supplemented diet and DNP is Donepezil.

In the prefrontal cortex, nitrite levels in the Scopolamine-only group were significantly elevated (4.22 ± 0.03) compared to the control group (3.13 ± 0.18), as shown in Figure 4B (F(5, 12) = 13.89; P < 0.0001). Treatment with OGSD at 5%, 10%, and 20%, along with Donepezil, significantly reduced nitrite levels to 3.04 ± 0.03, 3.24 ± 0.31, 2.62 ± 0.45, and 3.02 ± 0.20, respectively, compared to the Scopolamine-only group.

### Effects of *Ocimum gratissimum* on Catalase levels on the hippocampus and prefrontal cortex of scopolamine-induced memory impairment in Swiss mice

As shown in Figure 5A, Scopolamine administration significantly reduced (10.0 ± 2.00) catalase level in the hippocampus (F (5, 12) = 26.10; P<0.0001) when compared to the control (24.54 ± 2.01). Treatment with OGSD (5%, 10% and 20%) and Donepezil significantly increased catalase level (19.56 ± 1.93, 21.22 ± 1.34, 21.81 ± 1.79, and 20.91 ± 0.70) when compared with the Scopolamine-only group (10.0 ± 2.00).

**Figure 5.**
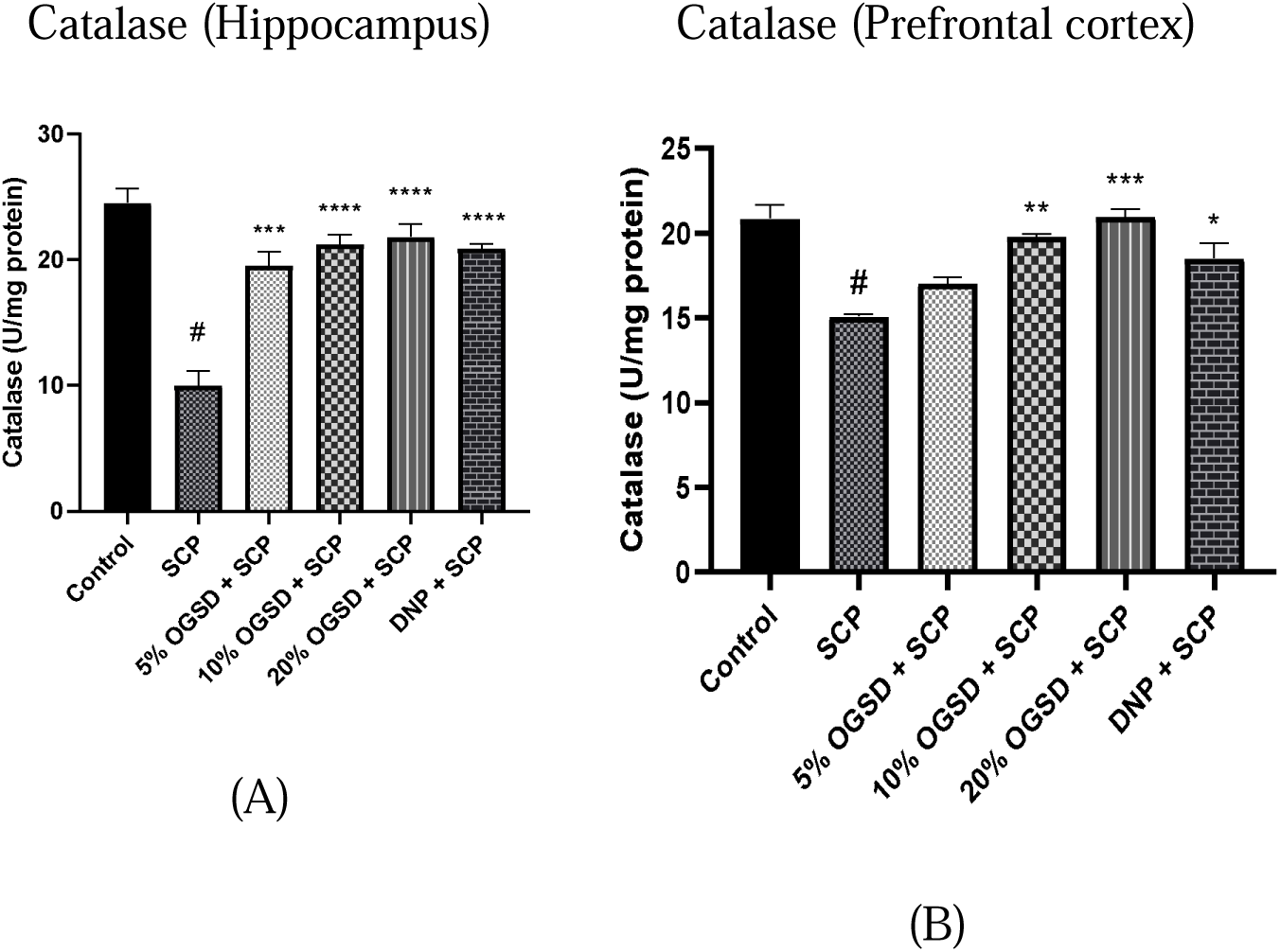
**A-B:** Effect of *Ocimum gratissimum*-supplemented diet on Catalase levels in Scopolamine-induced memory impairment. Values are represented as mean ± SEM (n=5). The data were analyzed using a One-Way Analysis of Variance (ANOVA) followed by a Sidak’s posthoc test. The significance level is expressed as # p < 0.05 for Scopolamine vs Control, * p < 0.05 for *Ocimum gratissimum* vs Scopolamine. SCP is Scopolamine, OGSD is *Ocimum gratissimum*-supplemented diet and DNP is Donepezil.

As shown in Figure 5B, Scopolamine administration significantly reduced (15.09 ± 0.22) catalase level in the prefrontal cortex (F (5, 12) = 17.10; P<0.0001) when compared to the control (20.86 ± 1.42). Treatment with OGSD (10% and 20%) and Donepezil significantly increased catalase level (19.80 ± 0.30, 20.96 ± 0.83, and 18.52 ± 1.53) when compared with the Scopolamine-only group (15.09 ± 0.22).

### Effects of *Ocimum gratissimum* on Superoxide Dismutase levels on the hippocampus and prefrontal cortex of scopolamine-induced memory impairment in Swiss mice

Scopolamine administration significantly reduced (0.87 ± 0.07) SOD level in the hippocampus (F (5, 12) = 30.85; P<0.0001) when compared to the control (1.71 ± 0.11) as shown in Figure 6A. Treatment with OGSD (20%) and Donepezil significantly increased SOD level (1.26 ± 0.01 and 1.29 ± 0.05) when compared with the Scopolamine-only group (0.87 ± 0.07).

**Figure 6.**
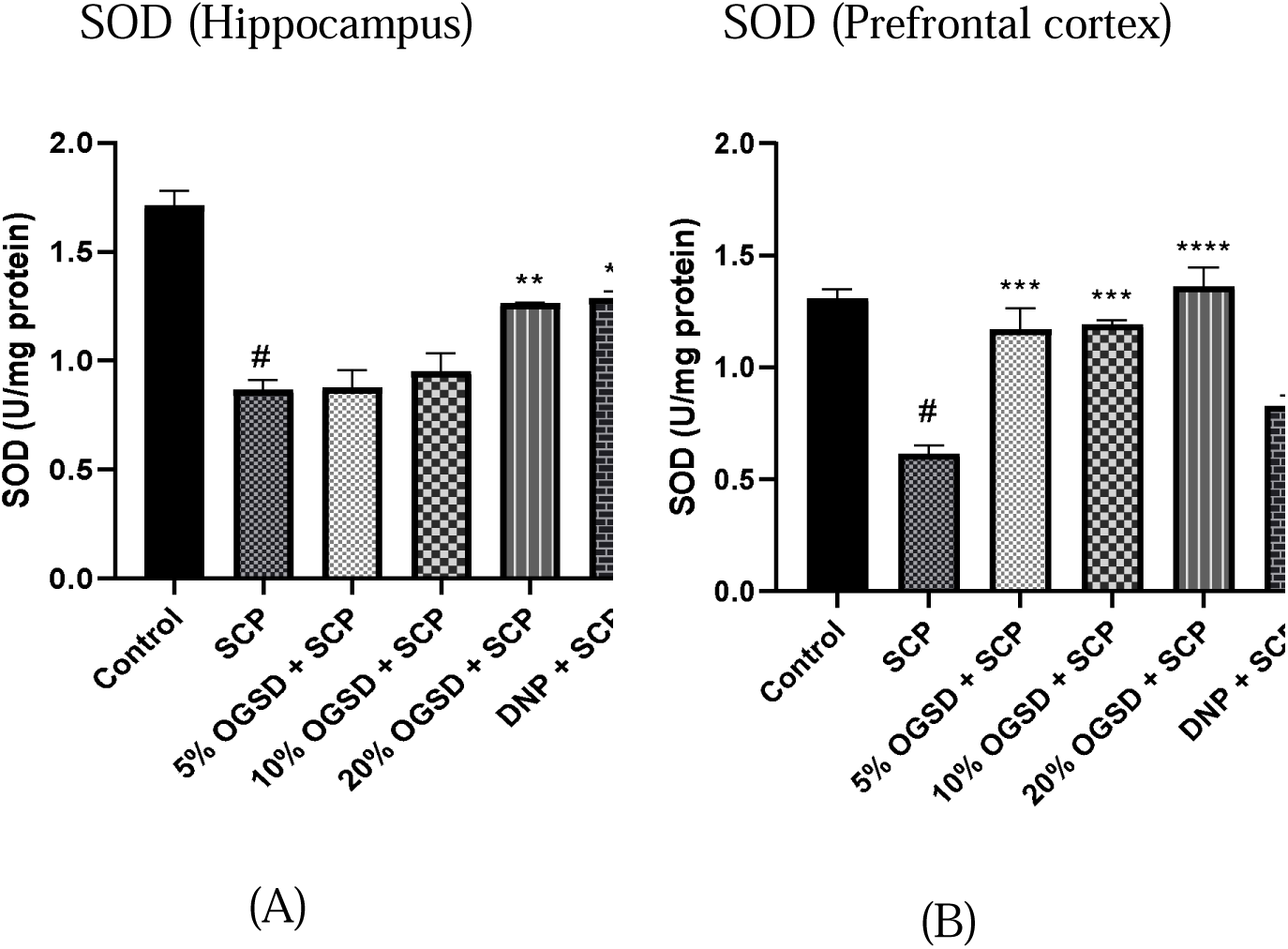
**A-B:** Effect of *Ocimum gratissimum*-supplemented diet on Superoxide Dismutase (SOD) levels in Scopolamine-induced memory impairment. Values are represented as mean ± SEM (n=5). The data were analyzed using a One-Way Analysis of Variance (ANOVA) followed by a Sidak’s posthoc test. The significance level is expressed as # p < 0.05 for Scopolamine vs Control, * p < 0.05 for *Ocimum gratissimum* vs Scopolamine. SCP is Scopolamine, OGSD is *Ocimum gratissimum*-supplemented diet and DNP is Donepezil.

Scopolamine administration significantly reduced (0.61 ± 0.07) SOD level in the prefrontal cortex (F (5, 12) = 23.1; P<0.0001) when compared to the control (1.31 ± 0.07) as shown in Figure 6B. Treatment with OGSD (5%, 10% and 20%) significantly increased SOD level (1.17 ± 0.17, 1.20 ± 0.04 and 1.36 ± 0.15) when compared with the Scopolamine-only group (0.61 ± 0.07).

### Effects of *Ocimum gratissimum* on Glutathione-S-Transferase levels on the hippocampus and prefrontal cortex of scopolamine-induced memory impairment in Swiss mice

Scopolamine administration significantly reduced (0.21 ± 0.01) SOD level in the hippocampus (F (5, 12) = 8.962; P<0.0001) when compared to the control (0.38 ± 0.05) as shown in Figure 7A. Treatment with OGSD (5%) significantly increased SOD level (0.34 ± 0.03) when compared with the Scopolamine-only group (0.21 ± 0.01).

**Figure 7.**
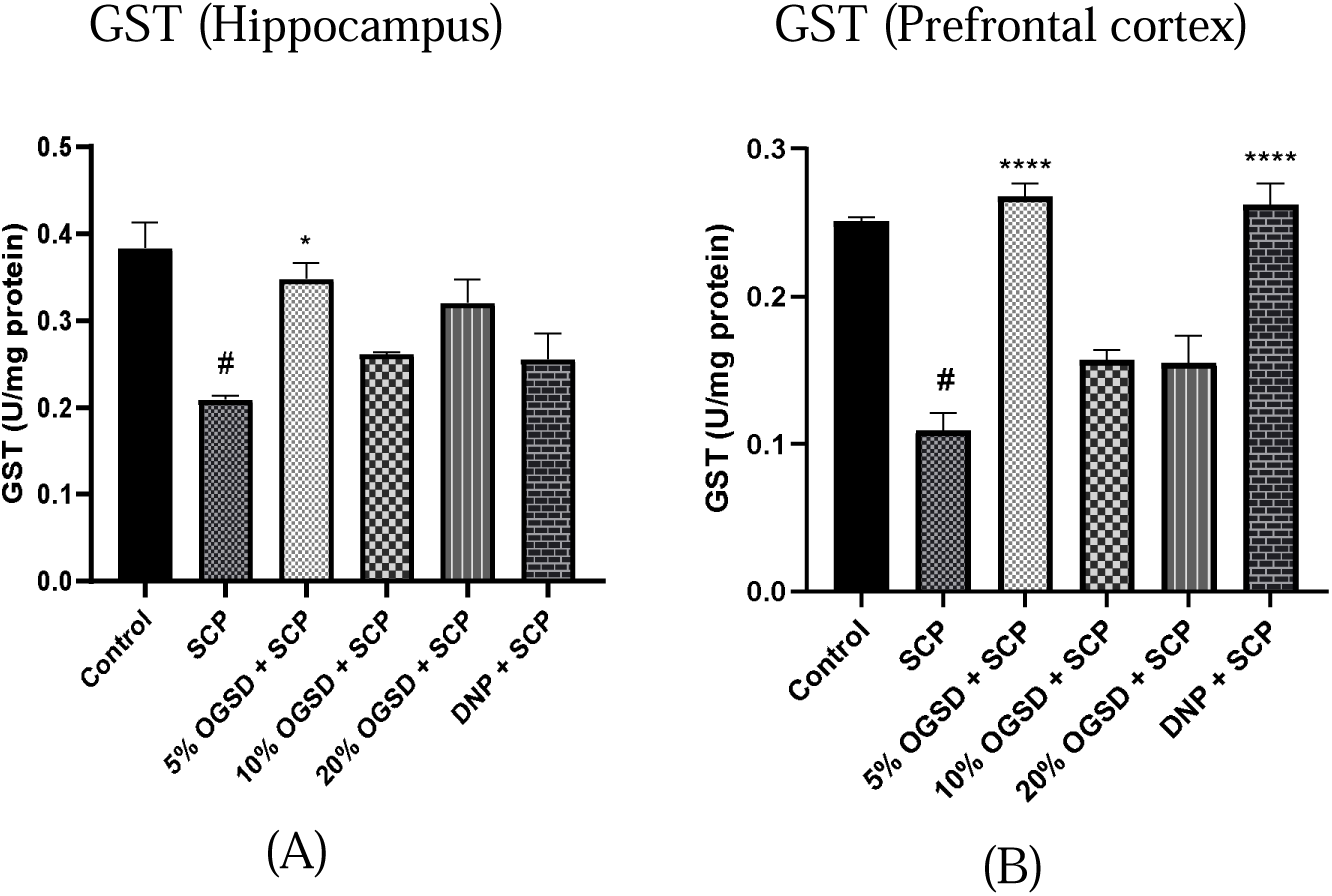
**A-B:** Effect of *Ocimum gratissimum*-supplemented diet on Glutathione-S-Transferase (GST) levels in Scopolamine-induced memory impairment. Values are represented as mean ± SEM (n=5). The data were analyzed using a One-Way Analysis of Variance (ANOVA) followed by a Sidak’s posthoc test. The significance level is expressed as # p < 0.05 for Scopolamine vs Control, * p < 0.05 for *Ocimum gratissimum* vs Scopolamine. SCP is Scopolamine, OGSD is *Ocimum gratissimum*-supplemented diet and DNP is Donepezil.

Scopolamine administration significantly reduced (0.11 ± 0.02) SOD level in the prefrontal cortex (F (5, 12) = 34.08; P<0.0001) when compared to the control (0.25 ± 0.004) as shown in Figure 7B. Treatment with OGSD (5%) and Donepezil significantly increased SOD level (0.27 ± 0.01 and 0.26 ± 0.02) when compared with the Scopolamine-only group (0.11 ± 0.02).

### Effects of *Ocimum gratissimum* on Proinflammatory cytokines TNF-α, IL-6 and acetylcholinesterase on the hippocampus and prefrontal cortex of scopolamine-induced memory impairment in Swiss mice

As shown in Figure 8A. TNF-α levels in the hippocampus (F (5, 12) = 18.70; P<0.0001) were significantly reduced in the OGSD-treated (20%) group (240.8 ± 32.13) and the Donepezil group (212.7 ± 22.07) compared to the Scopolamine-only group (359.1 ± 38.46).

**Figure 8.**
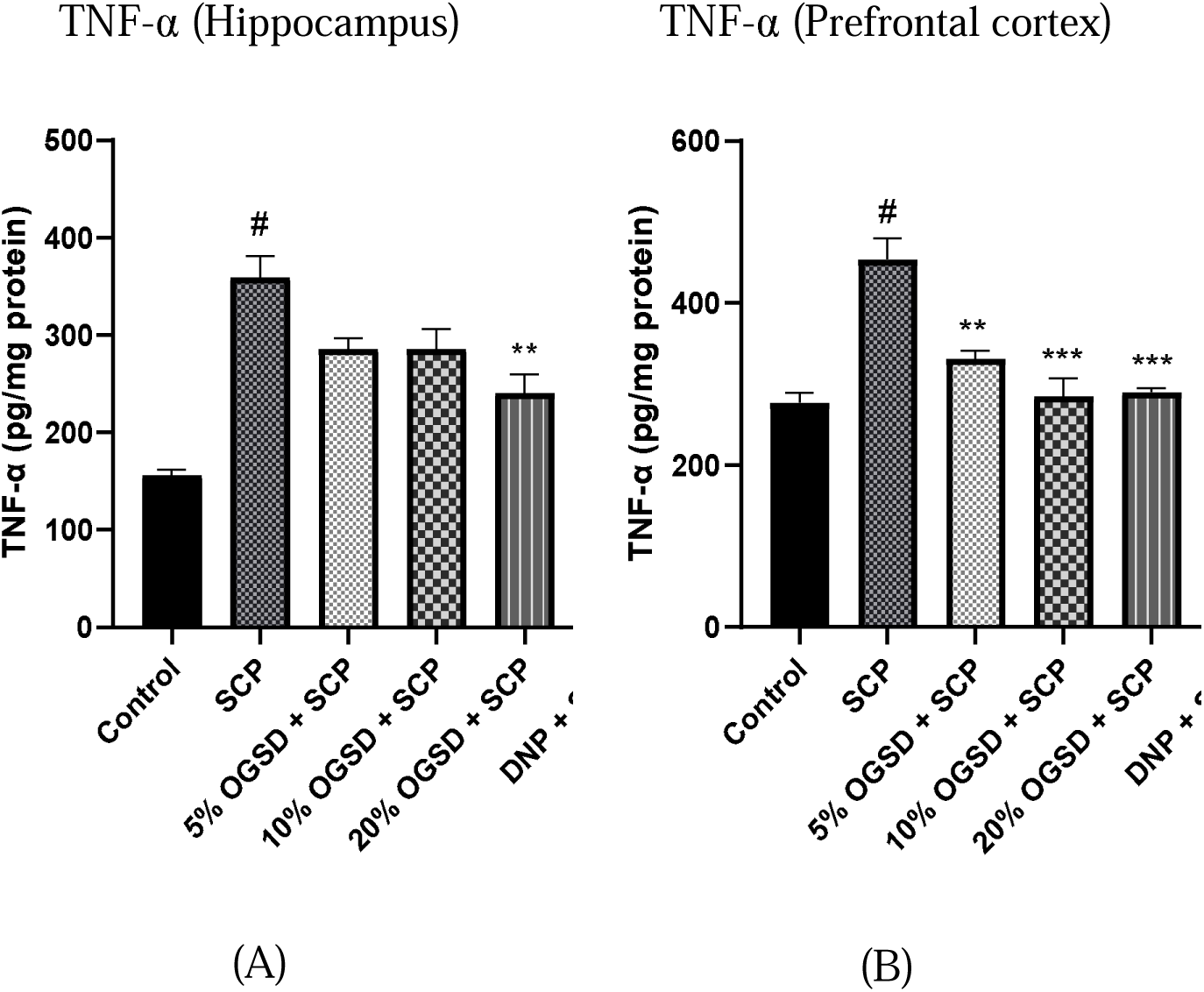
**A-B:** Effect of *Ocimum gratissimum*-supplemented diet on TNF-α levels in Scopolamine-induced memory impairment. Values are represented as mean ± SEM (n=5). The data were analyzed using a One-Way Analysis of Variance (ANOVA) followed by a Sidak’s posthoc test. The significance level is expressed as # p < 0.05 for Scopolamine vs Control, * p < 0.05 for *Ocimum gratissimum* vs Scopolamine. SCP is Scopolamine, OGSD is *Ocimum gratissimum*-supplemented diet and DNP is Donepezil.

**Figure 9.**
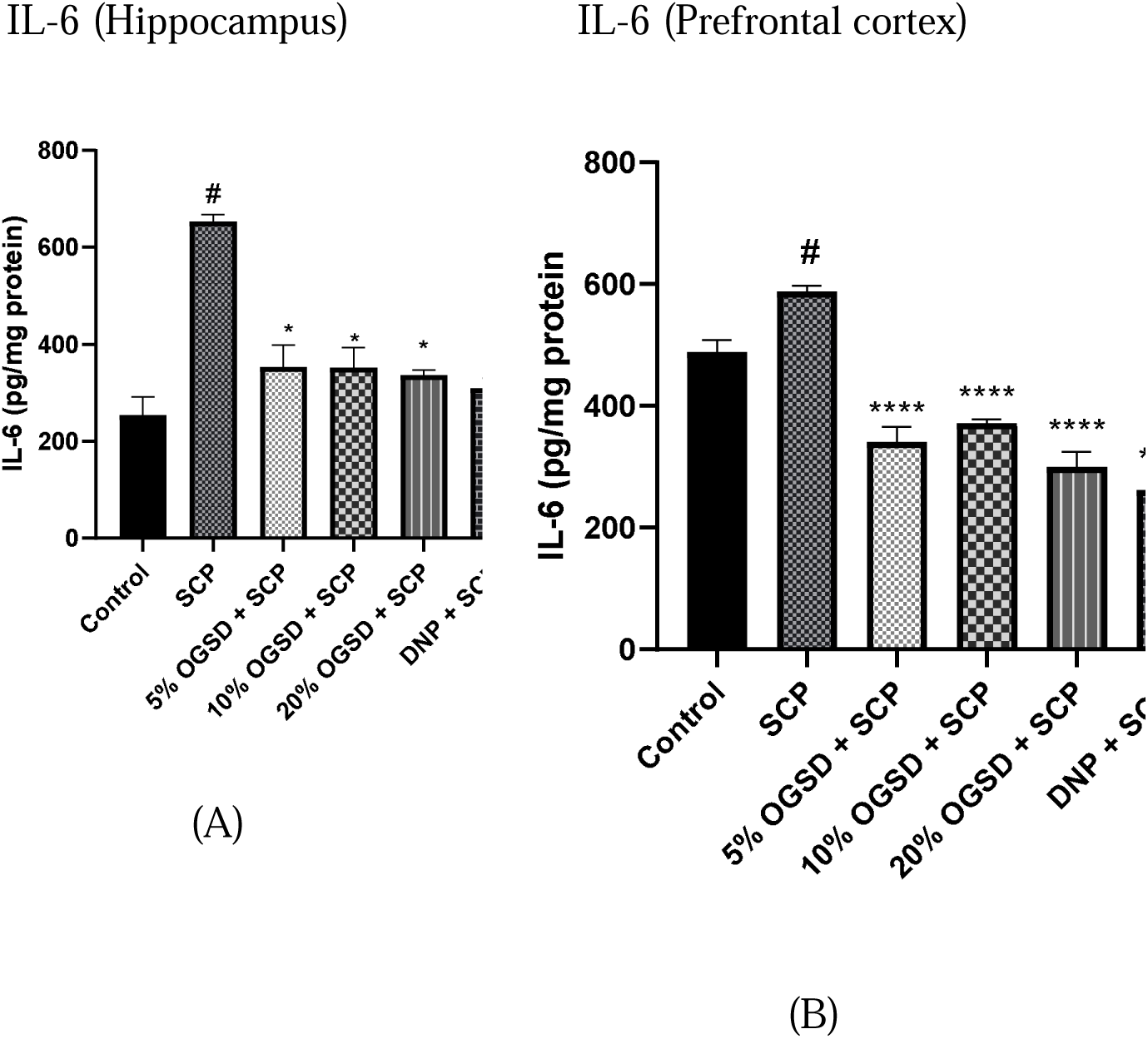
**A-B:** Effect of *Ocimum gratissimum*-supplemented diet on Interleukin-6 levels in Scopolamine-induced memory impairment. Values are represented as mean ± SEM (n=5). The data were analyzed using a One-Way Analysis of Variance (ANOVA) followed by a Sidak’s posthoc test. The significance level is expressed as # p < 0.05 for Scopolamine vs Control, * p < 0.05 for *Ocimum gratissimum* vs Scopolamine. SCP is Scopolamine, OGSD is *Ocimum gratissimum*-supplemented diet and DNP is Donepezil.

As shown in Figure 8B. TNF-α levels in the prefrontal cortex (F (5, 24) = 21.65; P<0.0001) were significantly reduced in all OGSD-treated groups (330.7 ± 17.50, 284.7 ± 39.35 and 289.6 ± 9.12) and the Donepezil group (216.4 ± 29.45) compared to the Scopolamine-only group (453.6 ± 45.72).

The Interleukin-6 (IL-6) levels (F (5, 12) = 20.98; P<0.0001) in the hippocampus were significantly reduced in all OGSD-treated groups (354.1 ± 75.74, 352.3 ± 70.92 and 336.5 ± 18.46) and the Donepezil-treated group (309.8 ± 35.57) compared to the Scopolamine-only group (653.4 ± 24.45).

The Interleukin-6 (IL-6) levels (F (5, 12) = 42.93; P<0.0001) in the prefrontal cortex were significantly reduced in all OGSD-treated groups (340.7 ± 42.06, 372.1 ± 9.57 and 300.0 ± 42.27) and the Donepezil-treated group (261.9 ± 36.13) compared to the Scopolamine-only group (587.7 ± 16.26).

Acetylcholinesterase activity in the hippocampus (F (5, 12) = 43.09; P<0.0001) were significantly reduced in all OGSD-treated groups (0.27 ± 0.02, 0.30 ± 0.01 and 0.36 ± 0.01) and Donepezil-treated group (0.18 ±0.03) compared to the Scopolamine-only group (0.42 ± 0.01). Acetylcholinesterase activity in the prefrontal cortex (F (5, 12) = 137.6; P<0.0001) were significantly reduced in OGSD-treated (20%) group (0.14 ±0.03) compared to the Scopolamine-only group (0.32 ± 0.02). Scopolamine administration also increased acetylcholinesterase activity as seen in the Scopolamine-only group (0.32 ± 0.02) compared to the control group (0.11 ± 0.01).

## DISCUSSION

In this study, two main well-established memory task models were used to assess memory in mice: Novel Object Recognition Test (NORT) and the Elevated Plus Maze (EPM) Test.

The effect of *Ocimum gratissimum*-supplemented diet (OGSD) on non-spatial recognition memory performance was assessed in mice using the Novel Object Recognition Test. The NORT measures non-spatial short and long-term memory and takes advantage of rodents’ unprompted nature to explore their natural surroundings. Therefore, NORT evaluates memory function based on the natural preference of rodents for novel objects (Tanglialatela, 2009). The discrimination index tends to quantify the ability to discriminate between a novel object and a familiar one and thus, evaluates memory function. The mice are typically discriminating between familiar and novel objects. The test is used in research to assess an animal’s ability to recognize and remember previously encountered objects. The lowest discrimination index in the scopolamine-only group demonstrated that scopolamine significantly impaired recognition memory on the Novel Object Recognition Test (NORT). However, mice given diets supplemented with 5%, 10%, and 20% *Ocimum gratissimum* exhibited markedly better discrimination indices, indicating a protective effect against memory impairment brought on by scopolamine. *O. gratissimum*’s neuroprotective and antioxidant qualities may be responsible for this improvement, as they most likely lessened cholinergic dysfunction. The result showed that OGSD caused a significant increase in the discrimination, which indicates that OGSD has neuroprotective potential on memory impairment (Ugbogu, 2009).

Spatial memory and navigation abilities were assessed in mice using the Elevated Plus Maze (EPM) Test by recording their transfer latency. The control group having the lowest transfer latency indicates that the animals have the shortest duration of time to transfer from one arm of the maze to another, which shows efficient spatial memory and navigation abilities (Figure 1B). The Scopolamine-only group has a higher transfer latency compared to the control group, and this implies that the animals took a longer time to transfer between maze arms, indicating that Scopolamine has impaired their memory and navigation ability. The OGSD-treated and standard drugs group showed a significant decrease in transfer latency, showing a mitigated Scopolamine-induced spatial memory impairment, which indicates the neuroprotective effect of *Ocimum gratissimum* on memory impairment.

Previous studies showed that Scopolamine increases Malondialdehyde level, indicating oxidative stress and lipid peroxidation (Khurana *et al*., 2016). It is formed as a byproduct when reactive oxygen species cause oxidative damage to cell membranes. The OGSD-treated groups had lower MDA levels compared to the Scopolamine-only group, indicating the neuroprotective effects of *Ocimum gratissimum*, leading to reduced lipid peroxidation. Glutathione is an antioxidant that tends to protect against oxidative stress by neutralizing reactive oxygen species (Dutta, 2020). Results obtained from this study showed that treatment with OGSD effectively restored the glutathione level, which was significantly suppressed by Scopolamine.

Nitrites are the stable products of nitric oxide metabolism. Elevated nitrite level signifies increased NO, which contributes to oxidative stress. The increased nitrite level in the Scopolamine-only group indicates excessive NO production (Rehman *et al*., 2017), contributing to the oxidative stress observed. Catalase tends to catalyse the decomposition of hydrogen peroxide to water and oxygen, thereby protecting the cells from oxidative damage. The low catalase level in the Scopolamine-only group compared to the control indicated indicated detoxification of hydrogen peroxide, leading to increased oxidative stress (Calderaro *et al*., 2022).

Superoxide dismutase is an important antioxidant enzyme that scavenges superoxide radicals (Ighodaro *et al*., 2018). This tends to defend against reactive oxygen species (ROS). The Scopolamine-only group having the lowest SOD level indicates reduced capacity to neutralize superoxide radicals thereby exacerbating oxidative stress. From the graph, the increased SOD level in the OGSD-treated shows a neuroprotective effect of *Ocimum gratissimum* on Scopolamine-induced memory impairment.

Glutathione S-transferase (GST) is an enzyme involved in detoxification processes, catalyzing the conjugation of GSH to various electrophilic compounds, making them more water-soluble and easier to excrete (Mannervik, 1985). The highest GST levels in the Scopolamine-only group reflect the compensatory response to detoxify the increased reactive species generated by Scopolamine.

Acetylcholinesterase and the cholinergic system are involved in learning and memory, and also play a major role in memory impairment. Studies have described a decrease in Acetylcholine and an increase in Acetylcholinesterase levels in the brain, contributing factors to memory dysfunction in neurodegenerative diseases like Alzheimer’s disease. In the result of this study, there was a significant increase in Acetylcholinesterase levels in the Prefrontal cortex and hippocampus in the Scopolamine-only group as compared to the control group (Figure 10A and 10B). However, in the prefrontal cortex, the Scopolamine group shows a high level of AChE activity compared to the control group. The administration of scopolamine must have led to an increase in AChE activity in the prefrontal cortex since it is known to impair cholinergic neurotransmission and has been linked to elevated AChE activity (Ghosh *et al*., 2019).

**Figure 10.**
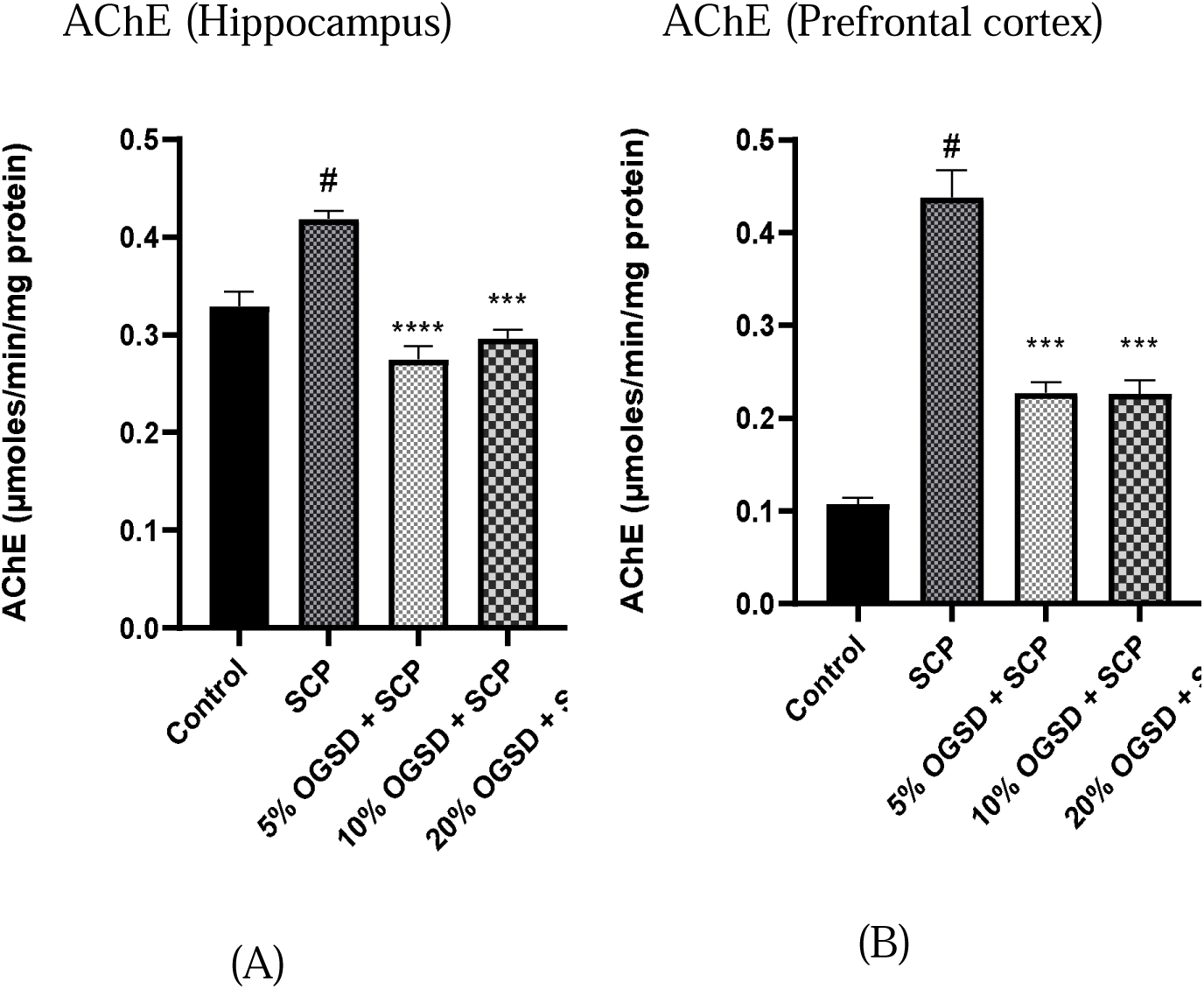
**A-B:** Effect of *Ocimum gratissimum*-supplemented diet on Acetylcholinesterase activity in Scopolamine-induced memory impairment. Values are represented as mean ± SEM (n=5). The data were analyzed using a One-Way Analysis of Variance (ANOVA) followed by a Sidak’s posthoc test. The significance level is expressed as # p < 0.05 for Scopolamine vs Control, * p < 0.05 for *Ocimum gratissimum* vs Scopolamine. SCP is Scopolamine, OGSD is *Ocimum gratissimum*-supplemented diet and DNP is Donepezil.

The Scopolamine-only group exhibited the highest levels of both TNF-α and IL-6 compared to the treatment groups. Tumor Necrosis Factor-alpha (TNF-α). It is a pro-inflammatory cytokine that plays a key role in the initiation and propagation of the inflammatory response. Elevated TNF-α levels have been associated with various neurological disorders, including cognitive impairment and Alzheimer’s disease. Similarly, interleukin-6 (IL-6) is another pro-inflammatory cytokine that has been implicated in neuroinflammation and cognitive dysfunction. Increased IL-6 levels as observed in the Scopolamine-only group, have been linked to the development of neuroinflammation and the subsequent impairment of cognitive processes.

In contrast, the OGSD-treated groups showed reduced levels of both TNF-α and IL-6 compared to the Scopolamine-only group. This suggests that OGSD intervention may have exerted anti-inflammatory effects, potentially mitigating the neuroinflammatory response induced by Scopolamine administration. Similarly, the Donepezil-treated group also exhibited a reduced level of TNF-α and IL-6 compared to the Scopolamine-only group. Donepezil, a commonly used cholinesterase inhibitor, has been shown to possess anti-inflammatory properties and may have contributed to the attenuation of the inflammatory response in this group.

This study investigated the neuroprotective potential of *Ocimum gratissimum*-supplemented diet on Scopolamine-induced Alzheimer’s disease-like memory impairment in Swiss mice. Scopolamine administration significantly impaired cognitive and performance memory in the Novel object recognition and Elevated Plus maze tests. Biochemical analyses revealed that scopolamine administration led to elevated levels of oxidative stress markers, such as MDA, NO, as well as increased pro-inflammatory cytokines (TNF-α and IL-6) in the hippocampus and prefrontal cortex which were mitigated by pretreatment with *Ocimum gratissimum*-supplemented diet.

## CONCLUSION

Firstly, it was observed that Scopolamine displayed impaired spatial memory and cognitive ability evidenced by the neurobehavioral and biochemical evaluations. However, different concentrations of *Ocimum gratissimum* shows promising results by improving the spatial memory and navigating abilities and also by improving the cholinergic transmission by inhibiting the action of acetylcholinesterase.

This study suggests that certain concentrations of *Ocimum gratissimum* have the potential to improve spatial memory and inhibit oxidative stress and acetylcholinesterase activity. These findings highlight the interplay between memory, oxidative stress, acetylcholinesterase activity and the effects of different treatment interventions.

It is recommended that further studies are needed to fully understand the mechanisms underlying these effects and to validate the efficacy of *Ocimum gratissimum* leaf as a potential therapeutic approach for cognitive impairment.

## DECLARATIONS

## Clinical trial number

not applicable.

## Ethics, Consent to Participate, and Consent to Publish declarations

The experimental protocol and sample size were in accordance with standard procedures and complied with the guidelines of the National Institutes of Health (NIH) for the care and use of laboratory animals. Ethical approval for this study was granted by the University of Ibadan Animal Care and Use Research Ethics Committee (UI-ACUREC) with approval number NHREC/UI-ACUREC/05/12/2022A.

Consent to participate and consent to publish are not applicable.

## Competing interest policy

The authors declare that they have no competing interest.

## Notes on contributors

**Damilare Alabi Akanbi** is an aspiring neuroscientist with a BSc. in Physiology from the University of Ibadan, Nigeria.

**Patrick Olanrewaju Ogboriga** holds a B.Sc. degree in Physiology from the University of Ibadan.

**Ismaheel Akinwale Adeniyi** is an aspiring neuroscientist. He holds a B.Tech and M.Sc degree in Physiology (Neuroscience). He is currently doing his PhD. at the Department of Physiology, University of Ibadan, Ibadan, Nigeria.

**Akintunde Muthoir Lawal** holds a B.Sc. and M.Sc degree in Physiology (Neuroscience). He completed his master’s degree in neurosciences at the Department of Physiology, University of Ibadan, Ibadan, Nigeria.

**Michael Ayomikun Olusola** holds a B.Sc. degree in Physiology from Ekiti State University

**Precious Ayomiposi Oluyemi** holds a B.Sc. degree in Physiology from Ekiti State University

**Samuel Adetunji Onasanwo** is a Professor, and Unit Head in Neurosciences and Oral Physiology, and the Head of the Department of Physiology, University of Ibadan, Nigeria.

## LIST OF ABBREVIATIONS

AChE: Acetylcholinesterase
AD: Alzheimer’s Disease
ANOVA: Analysis of Variance
BCA: Bicinchoninic Acid
CDNB: 1-Chloro-2,4-dinitrobenzene
CNS: Central Nervous System
DTNB: 5,5’-Dithiobis-(2-nitrobenzoic acid)
ELISA: Enzyme-Linked Immunosorbent Assay
EPM: Elevated Plus Maze
GSH: Glutathione
GST: Glutathione S-Transferase
IL-6: Interleukin-6
MDA: Malondialdehyde
NICE: National Institute for Health and Care Excellence
NO: Nitric Oxide
NORT: Novel Object Recognition Test
PBS: Phosphate Buffered Solution
ROS: Reactive Oxygen Species
SEM: Standard Error of the Mean
SOD: Superoxide Dismutase
TNF-α: Tumor Necrosis Factor-alpha

